# The recombination landscape of the barn owl, from families to populations

**DOI:** 10.1101/2024.04.11.589103

**Authors:** Alexandros Topaloudis, Eleonore Lavanchy, Tristan Cumer, Anne-Lyse Ducrest, Celine Simon, Ana Paula Machado, Nika Paposhvili, Alexandre Roulin, Jerome Goudet

## Abstract

Homologous recombination is a meiotic process that generates diversity along the genome and interacts with all evolutionary forces. Despite its importance, studies of recombination landscapes are lacking due to methodological limitations and a dearth of appropriate data. Linkage mapping based on familial data gives unbiased sex-specific broad-scale estimates of recombination while linkage disequilibrium (LD) based inference based on population data provides finer resolution data albeit depending on the effective population size and acting selective forces. In this study, we use a combination of these two methods, using a dataset of whole genome sequences and elucidate the recombination landscape for the Afro-European barn owl (*Tyto alba*). Linkage mapping allows us to refine the genome assembly to a chromosome-level quality. We find subtle differences in crossover placement between sexes that leads to differential effective shuffling of alleles. LD based estimates of recombination are concordant with family-based estimates and identify large variation in recombination rates within and among linkage groups. Larger chromosomes show variation in recombination rates while smaller chromosomes have a universally high rate which shapes the diversity landscape. We also identify local recombination hotspots in accordance with other studies in birds lacking the *PRDM9* gene. However these hotspots show very little evolutionary stability when compared among populations with shallow genetic differentiation. Overall, this comprehensive analysis enhances our understanding of recombination dynamics, genomic architecture, and sex-specific variation in the barn owl, contributing valuable insights to the broader field of avian genomics.

**Article summary:** To study recombination events we look either in family data or in population data, with each method having advantages over the other. In this study we use both approaches to quantify the barn owl recombination landscape. We find that differences exist between sexes, populations and chromosomes.

## Introduction

Recombination is a key feature of sexual reproduction that has multiple evolutionary implications but its inference is often overlooked in non-model species. Homologous meiotic recombination (hereafter recombination) is the reciprocal exchange of genetic material between homologous chromosomes during the first meiotic division. The physical exchange, called a crossover (CO), generates the necessary tension between chromosomes to ensure their proper segregation in the daughter cells and its absence has been associated with aneuploidy (Koehler *et al*. 1996; Hassold *et al*. 2007; Zickler and Kleckner 2015; Zelkowski *et al*. 2019).

Beyond contributing to the integrity of proper meiotic division, recombination has evolutionary consequences since it shuffles alleles between haplotypes thus affecting the genomic composition of a population. This shuffling can bring about evolutionary benefits including faster adaptation to a changing environment and more efficient selection (Hill and Robertson 1966; Otto and Lenormand 2002). However, recombination can also impede adaptation by breaking up beneficial combinations of alleles or increasing the rate of mutations and chromosomal rearrangements (Barton and Charlesworth 1998; Arbeithuber *et al*. 2015; Halldorsson *et al*. 2019). Furthermore, because recombination rates vary along the genomic sequence, they affect almost all genome-wide analyses. For instance, genetic diversity along the genome will correlate with recombination rates due to the effect of linked selection (Begun and Aquadro 1992) and regions of low recombination can appear as false positives in scans for selection based on differentiation (Booker *et al*. 2020). It is thus important to know the position of crossovers in the genome and the frequency with which they occur to account for recombination variation and eliminate confounding with other evolutionary forces.

However, quantifying recombination is a laborious task. One approach is linkage mapping, the positioning of markers along the sequence with a distance proportional to the recombination rate between them. This approach requires family data (or when available, controlled crosses) and has been applied to several species so far, providing a reliable measure of crossover frequency (Kong *et al*. 2002; Stapley *et al*. 2017; Brazier and Glémin 2022). Linkage mapping can also quantify the differences in recombination rates between sexes (i.e. heterochiasmy, Brekke et al., 2022; Johnston et al., 2017; Kong et al., 2010). Unfortunately, the family data required for linkage mapping is only available in certain species (Peñalba and Wolf 2020). In addition, recombination rates estimated with this method are limited by the number of meioses observed, making it impossible to quantify it accurately at a fine scale with the sample sizes available in most non-model species (Halldorsson *et al*. 2019).

To address this problem, additional approaches have been developed to estimate recombination using whole genome sequences from tens of unrelated individuals (Auton and McVean 2007; Chan *et al*. 2012; Spence and Song 2019). These methods model the observed linkage disequilibrium (LD) between markers, assessing ancestral recombination events that occurred in the coalescent history of the samples (Li and Stephens 2003; Stumpf and McVean 2003). This approach, hereafter referred to as LD-based inference, enabled the quantification of fine scale recombination variation in humans (McVean *et al*. 2004; Myers *et al*. 2005). In addition to human applications and because of the limited genomic resources required, LD-based inference of recombination has also been applied to non-model classes of species like birds, reptiles and fish among others (ex. Kawakami et al., 2017; Schield et al., 2020; Shanfelter et al., 2019; Singhal et al., 2015). These fine-scale inferences have identified the *PRDM9* gene to be essential for the location and rapid evolutionary turnover of recombination hotspots, narrow regions of increased recombination, in mammals (e.g. of mice and humans Booker et al., 2017; Myers et al., 2010). On the contrary, LD-based inference has shown that species that lack the *PRDM9* gene show either evolutionary conserved hotspots in regions of accessible chromatin, (e.g. dogs and zebra finches Auton et al., 2013; Singhal et al., 2015), or no hotspots at all (e.g. *C. elegans* and the genus *Drosophila* Kaur & Rockman, 2014; Smukowski Heil et al., 2015).

Despite its ability to quantify fine-scale variation with few genomes, LD-based inference has certain limitations. It infers the population recombination rate (rho), the product of the effective population size (N_e_) and the recombination rate, rather than the recombination rate directly. This has two major implications: i) LD-based inference does not distinguish between crossing over in male and female meioses and ii) is affected by forces that change N_e_ and not the recombination rate itself. The latter essentially means that forces that modify N_e_ (e.g. selection, demography) can confound estimates of recombination (O’Reilly *et al*. 2008). To remedy this and while selection remains a confounding factor, accounting for demography has been implemented in recent applications (Spence and Song 2019). Even then, estimates of LD-based methods are often validated with a different estimate of recombination, usually from other studies, such as linkage mapping (McVean *et al*. 2004; Axelsson *et al*. 2012; Shanfelter *et al*. 2019; Wall *et al*. 2022).

Therefore, a combination of approaches is the preferred route to accurately infer the recombination rates at the fine scale. However, such a combination requires both whole genome sequencing data of unrelated individuals and family data, a doubly costly resource. The availability of family data might be the most limiting factor but such resources are disproportionately available for certain classes of animals. For example, there is an over-representation of wild populations of birds because their nesting behaviour can facilitate population monitoring and can be exploited for the construction of long-term pedigrees (Grant & Grant, 2002; Lack & Lack, 1958; Pemberton, 2008). Despite this opportunity for the generation of family data in birds, we usually only have information about the recombination landscapes for a few genera. Most information for the order comes from studies of pedigreed populations using linkage mapping (Groenen *et al*. 2009; Backström *et al*. 2010; Kawakami *et al*. 2014; van Oers *et al*. 2014; Hagen *et al*. 2020; Peñalba *et al*. 2020; Robledo-Ruiz *et al*. 2022; McAuley *et al*. 2024) and we are aware of only two species where recombination was estimated both from an LD-based inference and a linkage-mapping approach: the zebra finch (*Taeniopygia guttata*) and the collared flycatcher (*Ficedula albicollis*) (Singhal *et al*. 2015; Kawakami *et al*. 2017).

All these previous studies have shown that recombination in birds exhibits broad-scale among-species variation in the absence of the *PRDM9* gene and inconclusive patterns of sex differences. Firstly, rates of recombination inferred from linkage mapping tend to differ between species despite a rather conserved avian karyotype (Ellegren 2010; Bravo *et al*. 2021). For example, two members of the passerine order (collared flycatcher and the superb fairy-wren *Malurus cyaneus*) show an unexplained twofold difference in recombination frequency (genetic length in cM) between their largest syntenic chromosome (Kawakami *et al*. 2014; Peñalba *et al*. 2020). On the contrary, through the loss of a functional *PRDM9* gene (Baker *et al*. 2017) birds show evolutionary stability of hotspots as demonstrated within finches and flycatchers (Singhal *et al*. 2015; Kawakami *et al*. 2017). It remains unknown how to reconcile the fine-scale stability of a recombination landscape without *PRDM9* with the broad variation observed among species. Further, birds show no consistent patterns of sex differences in recombination (heterochiasmy) with either males or females showing higher rates (Sardell and Kirkpatrick 2019; McAuley *et al*. 2024). However, until recently, conclusions on heterochiasmy were only based on the total recombination frequency (genetic length) summed over all chromosomes in each sex. In recent years, a few studies (Zhang *et al*. 2023; McAuley *et al*. 2024) have found evidence for differences in the placement of crossovers between sexes challenging the absence of sex differences in birds, when looking beyond the inconclusive linkage group analyses. Finally, since all avian species studied so far but the chicken (*Gallus gallus*) belong to the passerine order, it is hard to conclude if the sampled diversity is truly representative of the complete avian class.

Here, using both linkage mapping and LD-information, we present the first recombination landscape for a species of the owl order, the barn owl (*Tyto alba*). We use this species because it has the highest quality genome assembly for an owl species (Ducrest *et al*. 2020; Machado *et al*. 2022a), a set of whole genome sequences available from past studies (Machado *et al*. 2022a; b; Cumer *et al*. 2022a; b, 2024) and a long-term pedigreed population (Roulin 1999) with an untapped genomic potential (Charmantier *et al*. 2014; Sheldon *et al*. 2022). We capitalise on 176 genomes previously published along with 326 newly sequenced to build a high confidence variant set that spans the diversity of the species across the Western palearctic. For recombination inference, we use linkage mapping on a subset of our dataset, 250 owls belonging to 28 families to identify linkage groups in the barn owl sequence assembly, estimate the sex-averaged linkage map length and quantify sex differences in recombination. Additionally, we use an LD-based approach on 102 unrelated individuals from three populations to infer fine-scale recombination rate variation and scale our results using the estimates from the linkage map. With these complementary resources we quantify variation in recombination between sexes as well as identify substantial differences in fine scale patterns among chromosomes and populations.

## Results

In order to build the most comprehensive set of variants to date in the barn owl, we performed variant identification on 502 whole genome sequences of medium to high coverage (mean = 16, range = 8 to 43). Samples originate from 19 distinct localities spanning the Western Palearctic distribution of the species (with 3 to 13 samples from 19 localities, see Table S2 in File S2 for details). In addition to those, the Swiss population includes 346 individuals with a family structure originating from an observational pedigree. After filtering we retained 26,933,469 single nucleotide variants (SNPs) and used subsets of those individuals and variants for each analysis below (Table S4 in File S1).

### Linkage groups and recombination rate of the barn owl

After filtering the variant set on technical errors, allele frequency, distance and missingness (see Methods for more details), we ordered 154,706 variants along the 38 largest scaffolds of the genome assembly (Machado *et al*. 2022a) to create a linkage map for the barn owl. Based on segregation of these markers in 250 individuals from 28 families, we identified 39 linkage groups (LG). All the linkage groups identified correspond to scaffolds in the genome assembly except for Super-Scaffold 2 which was split into two linkage groups (see Supplementary text in File S1). The genome assembly of the barn owl therefore contains the sequence of 39 linkage groups out of 45 expected pairs of autosomal chromosomes (Table S1 in File S1). For these 39 linkage groups the final sex-averaged linkage map length spanned 2,066.81 centiMorgans (cM) over a physical sequence of 1,066 million base pairs (Mb) representing 88% of the genome assembly. Therefore, the genome average estimate of recombination rates for the barn owl is approximately 1.94 cM/Mb.

The genetic length of linkage groups increases only slightly with their physical length (Figure 1A). In general, each crossover that occurs per meiosis increases the genetic length of a linkage group by 50 cM (since two out of four products of the meiotic division are recombinant). The linkage groups of the barn owl showed an average genetic length of 53 cM (range: 25 – 83 cM) and all barn owl linkage groups recombine on average less than twice per meiosis (<100 cM). While the slope of the regression of the genetic length on the physical length is significantly positive (β = 0.417, 95% CI: 0.15 – 0.68), the intercept is less than the expected minimum of 50cM (α = 41.59, 95% CI: 33 – 50) under one obligate crossover per chromosome. In fact, a few linkage groups between 20 to 40 Mb of physical length have an inferred genetic length of less than 50 cM.

**Figure 1.**
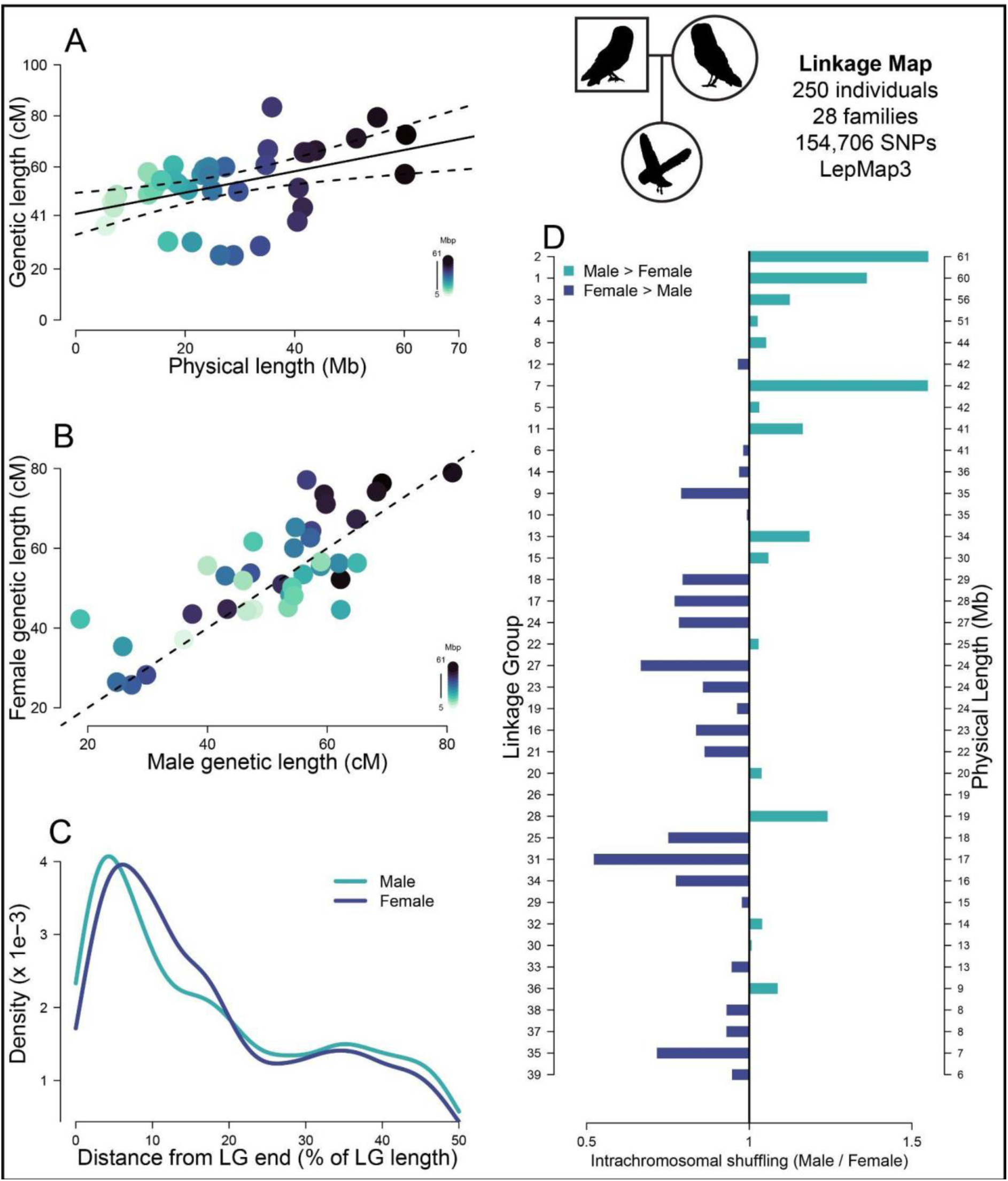
– A linkage map for the barn owl sheds light in fine scale heterochiasmy. All plots in this figure are products of the linkage map dataset consisting of 250 individuals in 28 families, illustrated with the pedigree of owl symbols on the top right. **A:** Linkage mapping estimates of sex-averaged genetic lengths for the linkage groups identified in the barn owl assembly plotted against their physical length. Regression line is shown with 95% confidence intervals (α=41.6, β=0.4171, t=3.093, p=0.004). Colour intensity scales with linkage group physical lengths as in legend. **B**: Recombination map length (cM) of linkage groups (LG) for females plotted against the recombination map length for males. Dashed line is the identity (y=x) line. Each dot represents one linkage group and the colour intensity scales with their physical lengths. **C**: Density plot of male (aquamarine) and female (blue) crossover (CO) counts plotted along the distance from the LGs end. X-axis is in percentages of total linkage group sequence. Density values are scaled so that they sum to 1. **D:** Differences between sexes in rates of intrachromosomal shuffling (r_intra) presented as a ratio (male r_intra / female r_intra) for different LGs. Bars to the right of the black line coloured green signify higher intra-chromosomal shuffling in males, while bars to the left of the black line coloured in blue correspond to linkage groups with higher shuffling in females. LGs are ordered by decreasing physical length (larger LG on top) as signified on the second y-axis on the right.

### Subtle heterochiasmy

To infer heterochiasmy, we look at the sex-specific linkage map estimates (Figure S6 & Table s1 in File S1). Female barn owls, with a map length of 2,124 cM, have a 5% larger genetic map than male barn owls (2,013 cM). There appears to be no consistent pattern of heterochiasmy among linkage groups (Figure 1B). To investigate potential differences in localisation of recombination events in males and females, we look at the positioning of crossovers along the length of all linkage groups (Figure 1C). Overall in the barn owl, crossovers tend to occur closer to the linkage group extremities than in their centre. However, the distribution of crossovers differs between the sexes (Two-sample Kolmogorov-Smirnoff test D=0.197; p < 0.001). Notably, males tend to recombine more at the extremities and the middle of the linkage groups.

Because sexes showed different locations of crossovers and not all crossovers are as effective at shuffling alleles between haplotypes, we quantified the rate of intra-chromosomal shuffling (r_intra) as defined in Veller et al. (2019). Briefly, this quantity measures the relative shuffling of alleles due to a crossover along the length of the chromosome. For example, a crossover in the middle of the chromosome shuffles more alleles than a distal one. We estimated rates of intra-chromosomal shuffling in males and females (Figure 1D). Despite an overall lower recombination frequency (Figure 1B) males show up to 50% higher intra-chromosomal shuffling for larger linkage groups (Figure 1D). On the other hand, females show higher rates of shuffling in intermediate to smaller linkage groups.

### Fine scale variation among linkage groups

To investigate finer scale variation in recombination rates, we turn to recombination rates estimated from patterns of linkage disequilibrium (LD) (Figure S5 in File S1). We estimated recombination rates using pyrho in a set of 9.3 million variants (number of variants per population are presented in Table S4 in File S1) identified along the whole genome of 76 unrelated birds from Switzerland (CH). The total genetic length estimated from LD was 957 cM, 2.1 times less than the linkage map estimate for the same population. We scaled the total length inferred from LD to be the same as the linkage mapping estimate, to account for the confounding effect of N_e_ and compared the estimates in non-overlapping 1 Mb windows. The correlation of recombination rate estimated from the linkage map and from pyrho at the 1 Mb scale was high (r = 0.88, 95% CI: 0.869 – 0.896) but pyrho showed higher estimates in regions of low recombination compared to linkage mapping (Figure 2A).

**Figure 2.**
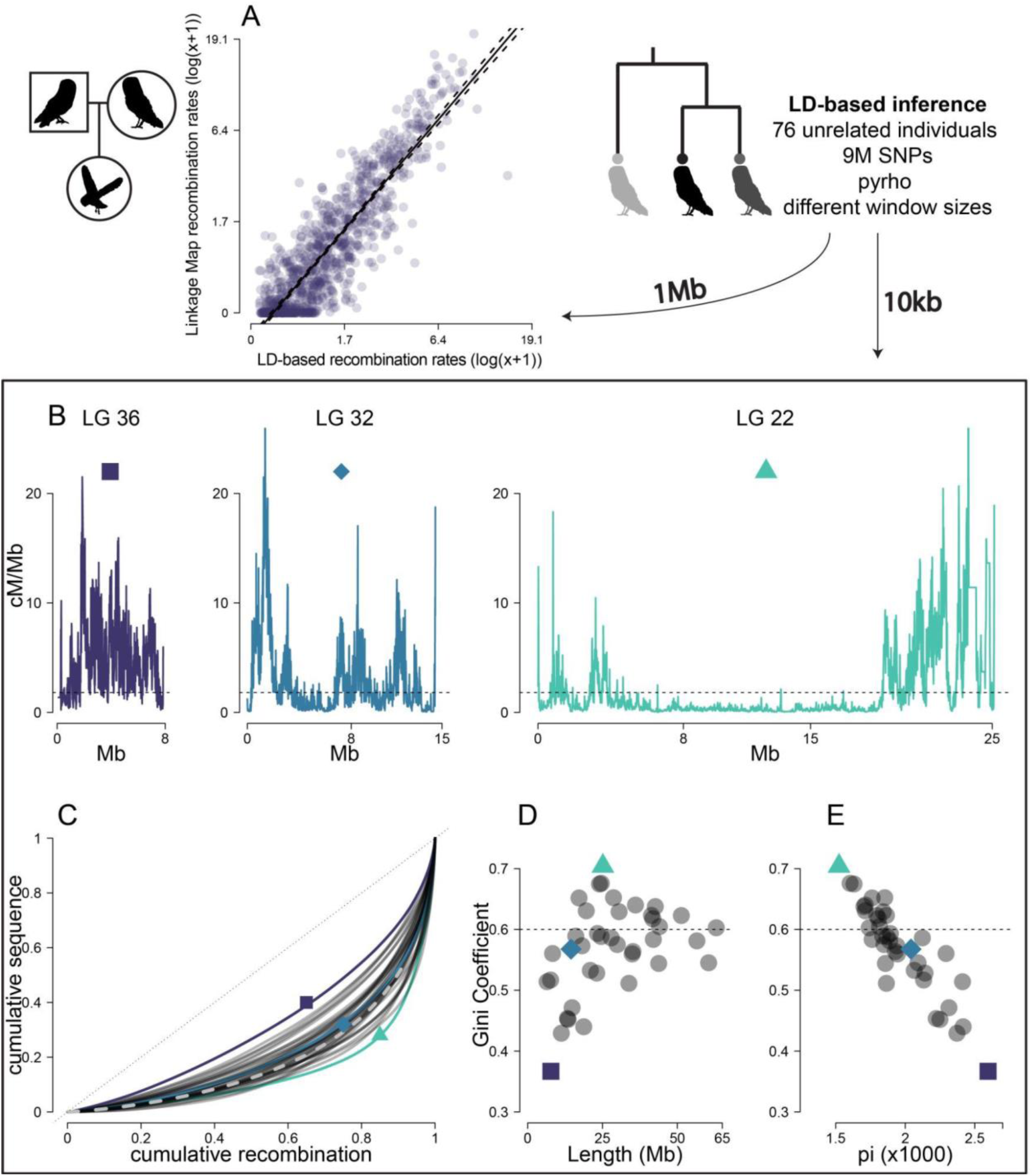
– Variation of recombination among linkage groups. As in figure 1, pedigree of owls symbol signifies results from linkage mapping. On this figure and later figures the tree of owls shows results from pyrho along with the window resolution used next to it or above it, here 1 Mb and 10 kb. **A:** Comparison of recombination rate estimates from linkage mapping and LD inference for the Swiss population. The comparison is made in 1 Mb windows to avoid inaccurate linkage mapping estimates due to the limited number of meioses observed. Axes are in the natural logarithm of the value + 1 to limit values to positive range. Regression line is shown with prediction intervals as dashed lines (α=-0.285, β=1.2, t=62.92, p<0.001) **B:** The recombination frequency (cM/Mb) in 10 kb windows along the physical map of three example linkage groups (LG 36, LG 32 and LG 22 respectively). The dashed horizontal line is the genome average recombination rate. These linkage groups represent linkage groups with different recombination landscapes. The purple square linkage group (LG 36) has the most evenly spread recombination along its length, the blue diamond (LG 32) has an intermediate spread that corresponds to the genome average and the aquamarine triangle (LG 22) has the most punctuated landscape. **C:** Cumulative sequence plotted against cumulative ordered recombination length for each linkage group (dark grey curves) and genome-wide (grey dashed curve). The black dotted line is the identity (y=x) line. The Gini coefficient corresponds to the area delimited by each curve and the identity line. **D**: The Gini coefficient of recombination rates for each linkage group plotted against its physical length. Dashed grey line is the genome average Gini coefficient. **E**: The Gini coefficient of recombination plotted against the average nucleotide diversity of each linkage group. Dashed grey line is the genome average Gini coefficient.

The recombination landscape differed among chromosomes with different sizes. To further quantify this variation in recombination rates, we quantified the proportion of genomic sequence where recombination occurs. By ordering all 10 kb windows for each linkage group in decreasing recombination rates, we plot the cumulative recombination percentage against the cumulative percentage of sequence (Figure 2C). Overall, 80% of recombination occurs in approximately 35% of the sequence (dashed grey line in Figure 2C). However, there is substantial variation in the distribution of recombination among linkage groups. To further measure this skewness, we used the Gini coefficient of recombination rates for each chromosome. Briefly, the Gini coefficient corresponds to the area between each curve in Figure 2C and the identity (y=x) line and ranges from 0 to 1. Smaller values indicate an evenly spread landscape (every window has the same recombination rate) and higher values a more rugged one (windows show large differences in recombination rates). The genome-wide average Gini coefficient is 0.6 and the linkage group specific estimates varied between 0.37 and 0.70 (LG 36 and LG 22, respectively marked with a blue square and a green triangle in Figure 2). Along with a linkage group of an intermediate Gini coefficient (0.57, LG 32 – aquamarine diamond in Figure 2), their recombination landscapes are presented in Figure 2B. We found that the Gini coefficient depends on the physical length of the linkage group with more evenly spread (and elevated) recombination rates in smaller linkage groups and more concentrated landscapes in larger ones (Figure 2D) but the effect seems to reach a plateau as the length increases above 25 Mb. Further, the Gini coefficients correlate negatively and strongly with the average nucleotide diversity of the linkage group with more concentrated recombination peaks leading to lower average nucleotide diversity (Pearson’s r = –0.9, 95% CI: –0.947, –0.82) (Figure 2E). Overall, recombination rates vary substantially among the different linkage groups of the barn owl assembly.

### Identifying hotspots of recombination

Because birds lack the *PRDM9* gene, recombination hotspots are expected to localise to transcription start and end sites (TSS, TES respectively), as well as CpG islands (CGIs) (Singhal *et al*. 2015; Baker *et al*. 2017). To verify this, we used estimates of recombination frequency in non-overlapping windows of 1,000 base pairs (1 kb) along the genome. Windows that were annotated to contain either a TSS or a TES (n=30,224, 2.7% of windows) or contained a CGI spanning the whole window (n=13,841, 1.2% of windows) were identified and their recombination rate was divided by the average recombination rate in 40 kb upstream and 40 kb downstream of the focal window (relative recombination rate in 80 kb – RRR80). The results showed elevated recombination rates in the focal windows compared to their vicinity (Figure 3A).

**Figure 3.**
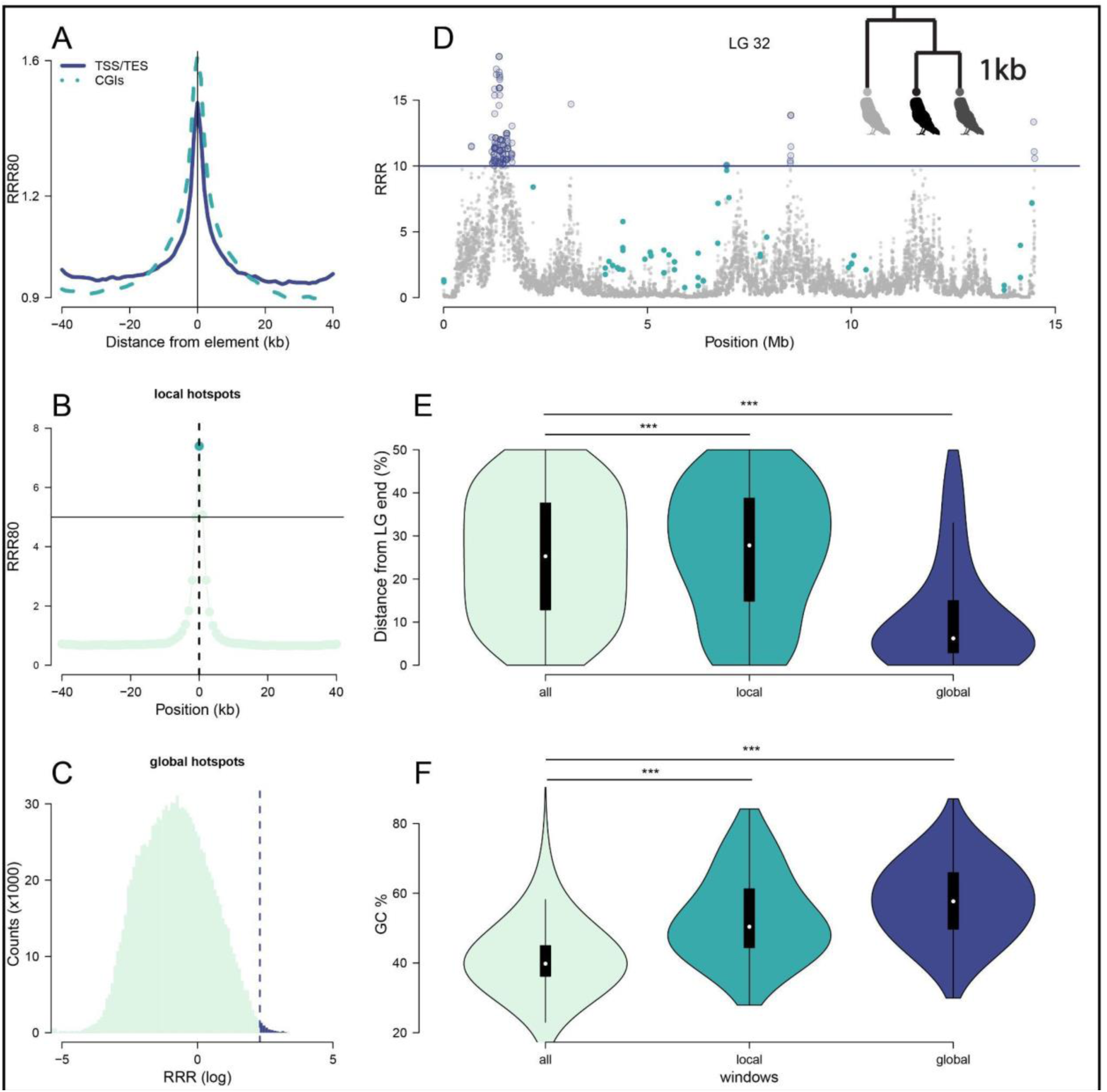
Hotspot characteristics of the barn owl in 1 kb windows. All results on this figure were obtained using pyrho in 1 kb windows as illustrated with the tree of owls. **A:** Lacking *PRDM9*, recombination is increased at the local regions around annotated transcription start and end sites (TSS,TES, full line) as well as CpG islands (CGIs, dashed line). RRR80 is the recombination rate of each 1 kb window divided by the average in 80 kb around. The lines show the average across all identified elements. **B:** Local hotspots are defined as 1 kb windows with a recombination rate at least five times higher than the average in 80 kb around (RRR80 > 5). This example (which is an average plot of all windows annotated as local hotspots) the focal window (aquamarine dot at position 0) would be annotated as a local hotspot. **C:** Global hotspots are defined as 1 kb windows with a recombination rate at least ten times the genome-wide average (RRR > 10). All windows right of the dashed line are global hotspots. The RRR is given in log scale for better representation **D:** Example of annotated global and local windows in linkage group (LG) 32. Aquamarine dots are local hotspots while the purple line shows ten times the global average and thus all windows above the line are global hotspots as signified with the open purple circles. **E:** Violin plot of the location of all windows, local and global hotspots relative to the end of the linkage group. Local hotspots are usually found towards the middle of the linkage groups because they are identified in regions of lower background recombination. On the other hand global hotspots are concentrated around the ends (as expected from Figure 1C). Significance assigned through a Wilcoxon rank sum test. **F:** GC content is elevated in both local and global hotspots. Significance assigned through a Wilcoxon rank sum test.

The inequality in the distribution of recombination rates along the sequence presented in Figure 2C supports the existence of recombination hotspots in the barn owl (Myers *et al*. 2005). We thus looked for recombination hotspots at the kilobase resolution. In species without *PRDM9*, past studies focus on local hotspots. We define local hotspots as 1 kb windows that exhibit five times the average recombination rate in 80 kb around the focal window (RRR80 ≥ 5, Figure 3B). Since the definition of a local hotspot is arbitrary and cutoff levels differ (e.g. Myers et al., 2005, Singhal et al., 2015, Kawakami et al., 2017), we chose to follow Singhal et al., 2015. To compare local hotspots against a more robust set of hotspots we also annotated global hotspots following (Halldorsson *et al*. 2019) as 1kb windows that show a recombination rate higher than ten times the genome average (RRR ≥ 10 – Figure 3C).

In the Swiss population, we identified a total of 3,949 local hotspots containing 1.8% of the total genetic length and 4,440 global hotspots containing 5.5%. 499 windows were annotated as both local and global hotspots. Local hotspots were usually identified in regions of lower recombination rates which are found towards the middle of linkage groups (Figure 1C) while global hotspots were by definition in peaks of recombination, concentrated around the ends (example in Figure 3D, Figure 3E). Both hotspot classes showed higher GC content distribution compared to the genome average (Figure 3F) which supports their annotation as hotspots. GC content was higher in global hotspots than local ones.

### Low repeatability of fine-scale recombination landscapes

To quantify the change of the recombination landscape across European populations of the species, we used three populations from the Western Palearctic, Portugal (PT, n=13), Great Britain (GB, n=13), and Switzerland (CH, n=76) (Figure 4A). For Portugal we pooled together three samples from Morocco and ten from Portugal since they have very high genetic similarity (Cumer *et al*. 2022a). Because the sample size in Switzerland was far larger than the other two populations and to test the robustness of our results, we randomly subsampled two sets of 13 individuals from Switzerland, creating pseudo-replicate populations (CH13, CH13_2). For Portugal and Great Britain genetic length estimates were 1,271 and 1,345 cM respectively. These estimates closely matched those for the subsampled sets from Switzerland which were 1,327 and 1,296 cM, respectively. For all populations, we scaled the results so that the total genetic length would match that of the sex-averaged linkage map estimate of 2,067 cM. After scaling, we compared all populations with the linkage map in 1 Mb windows along the genome and found that genome-average correlations were higher than 0.8 (Figure S1 in File S1). At finer scale, however, recombination landscapes varied among populations (an example landscape for all populations is presented at the 1 kb scale for the first five Mb of LG 32 in Figure 4B). Estimates in GB showed reduced resolution at the finer scales (aquamarine line in Figure 4B), probably because of reduced genomic diversity and historical effective population size in that population (Table S3 & Figure S2 in File S1, see also Machado, Cumer, et al., 2022).

**Figure 4.**
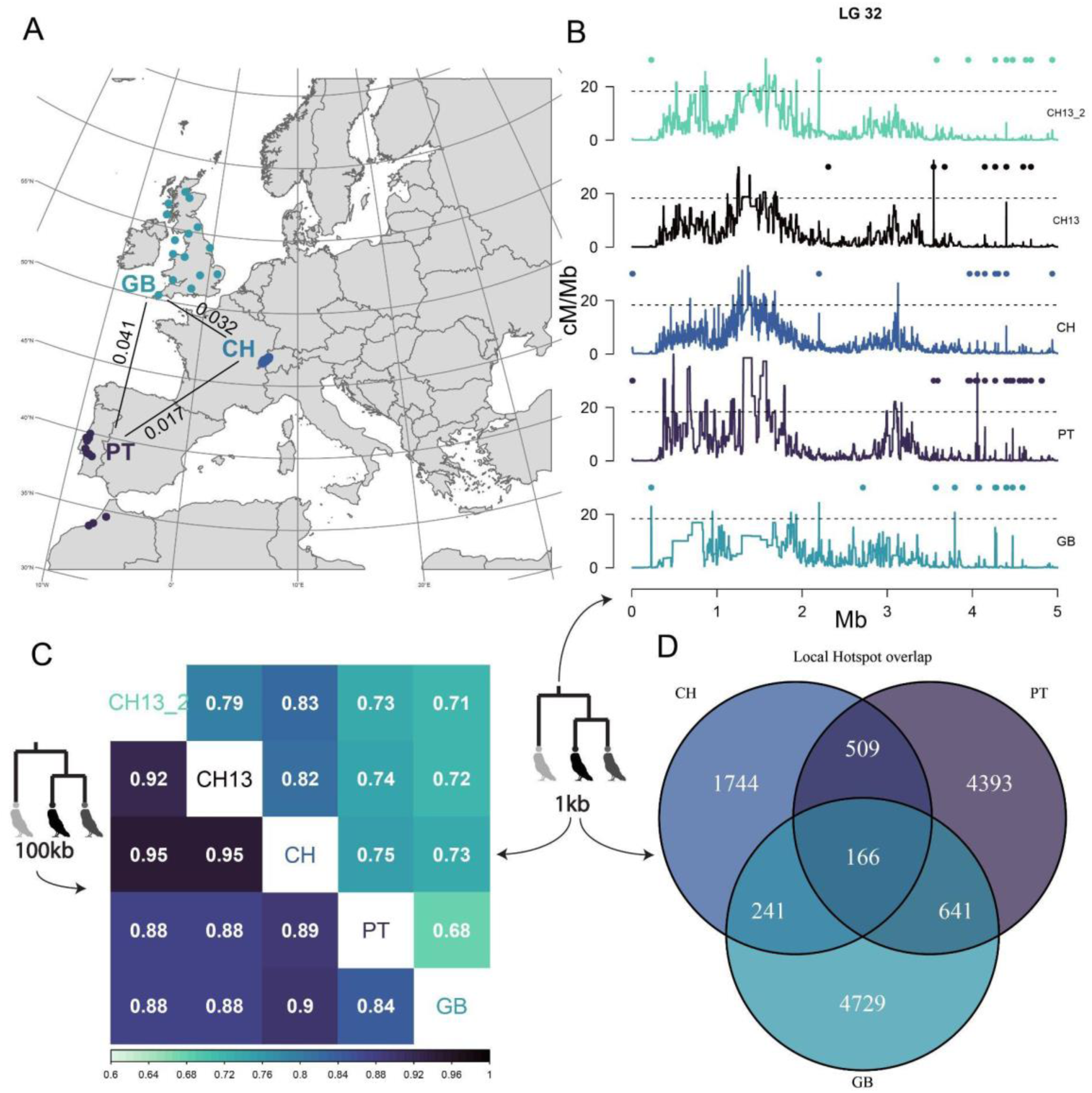
Comparison of recombination landscapes between populations. **A**: Map of sampled populations. Numbers correspond to genome-wide pairwise F_ST_ values (Machado *et al*. 2022a) **B:** Example of recombination landscape for 1 kb windows in all datasets for the first 5Mb of linkage group (LG) 32. Points above each plot show local hotspots for each population. Local hotspots are identified as in Figure 3 (RRR80 >5). The dashed line is the line above which we defined global hotspots (10 times the average for each population). **C:** Correlation matrix of recombination rates for 1 kb windows (above diagonal) or 100 kb windows (below diagonal). **D:** Venn diagram of local hotspot counts within and between populations. Population codes are as follows: CH: full Swiss dataset (n=76), PT: Portuguese dataset (n=13), GB: Great Britain dataset (n=13), CH13: undersampled first Swiss dataset (n=13), CH13_2: undersampled second Swiss dataset (n=13).

We quantified the divergence of the recombination landscapes as the correlation of recombination rates on different window sizes (1 kb and 100 kb) among all pairs of populations (Figure 4C). Two main patterns emerge from the comparison of population landscapes. First, correlation of landscapes depended on the scale used, with narrower windows showing smaller absolute values (above and below diagonal in Figure 4C). Second, in both scales, correlations of CH with PT and GB were larger than correlations of PT and GB with any subsampled Swiss datasets (CH13, CH13_2). However, as expected the Swiss datasets (CH, CH13, CH13_2) resembled one another more closely (Figure 4C and Figure S3 in File S1). Concerning hotspots, datasets with smaller sample sizes exhibited more local hotspots than the full Swiss dataset (Figure 4D). At the same time, local hotspot overlap was poor (less than 50%) between any pair and among all three populations (Figure 4D & Figure S4 in File S1). This pattern was replicated even when comparing hotspot sharing among the Swiss datasets (Figure S4 in File S1). In fact, CH and CH13 shared a lower percentage of local hotspots than did PT and CH. In general, global hotspots showed higher values of sharing than local hotspots and more consistent patterns of sharing (Figure S4 in File S1).

## Discussion

Recombination, the shuffling of alleles during meiotic division, is a major mediator of evolution but we know little about the recombination landscape of most species. In this study, using an extensive whole genome sequencing (WGS) dataset of barn owl, we infer recombination with two different methods to describe broad and fine scales of recombination variation: linkage mapping on a pedigreed population and an linkage-disequilibrium(LD) based approach on three different populations. We identify linkage groups in the barn owl genome assembly and quantify the recombination rate in the species to be approximately 2cM/Mb. We show that in the barn owl, overall length of genetic maps are not very different between males and females but sexes show subtle, fine-scale differences in crossover placement and shuffling proportions. Despite few (1-2) crossovers per chromosome, we find large variation in the location of these crossovers among different linkage groups. We show that this variation shapes the diversity landscape and is only partially determined by the linkage group’s physical length. Looking at the kilobase scale, recombination rates are increased in windows that contain transcription start and end sites and CpG islands, as is expected from a species without *PRMD9* (Baker *et al*. 2017). At the same time, local hotspots are found in regions of lower average recombination (usually at the centre of linkage groups) and show an elevated GC ratio, but still lower than that of global hotspots. Lastly, population comparisons show local recombination hotspots locations vary between populations despite low genetic differentiation and high broad-scale recombination correlation. We discuss these results and their implications below.

### Linkage groups confirm the near completeness of the barn owl assembly

Complete genome assemblies are a precious resource that requires multiple sources of information. Beyond read acquisition, assembling a genome requires an ordering process to orient the reads into scaffolds and the scaffolds into chromosomes. In-silico this can be achieved using physical mapping of the reads (Lieberman-Aiden *et al*. 2009; Burton *et al*. 2013), or through the use of linkage mapping (Fierst 2015) although linking computationally assembled scaffolds to karyotypic chromosomes will eventually require molecular techniques such as FISH for confirmation (Shakoori, 2017). The latest barn owl assembly (Machado *et al*. 2022a) was assembled into super-scaffolds using optical genome mapping (BioNano, Lam *et al*. 2012). In the present study, we verified and improved the barn owl assembly by anchoring the largest 38 scaffolds into 39 linkage groups and revealed that the genome assembly of the barn owl is of chromosome-level quality.

The karyotype of the barn owl contains 45 autosomal pairs (Belterman and De Boer 1984; Peona *et al*. 2018). Therefore, the linkage map is still missing 6 autosomes. We note that these elements might be partially present in the physical assembly since smaller scaffolds with a few tens of identified markers that passed filtering could not be confidently allocated to linkage groups. Regardless, the chromosomes missing are probably the smallest six microchromosomes, or dot chromosomes, notoriously difficult to sequence and assemble due to their high GC content and reduced chromatin accessibility (Burt 2002; Bravo *et al*. 2021; Waters *et al*. 2021). Notably, these microchromosomes were only recently assembled in the chicken genome (Huang *et al*. 2023) and are missing from most available bird reference genomes (Peona *et al*. 2018). In fact, most microchromosomes will remain elusive until future studies make use of advances in long-read technologies (Marx 2023) to complete the reference genomes of birds. In this endeavour, linkage mapping, when available, can be an invaluable tool (e.g. Peñalba et al., 2020; Robledo-Ruiz et al., 2022).

### Differential shuffling between sexes due to crossover placement

Our results point at a fine scale variation in crossover placement between sexes that is not immediately apparent when investigating sex differences at the linkage group scale. Furthermore, this variation leads to a differential shuffling of markers in each sex among different chromosomes. While in mammals, heterochiasmy seems to point to increased rates in females especially around centromeres, in avian studies results are inconclusive on a general pattern of heterochiasmy in the class. For example, male collared flycatchers exhibit higher genetic lengths than females while in a recent study of the great reed warbler (*Acrocephalus arundinaceus*) no differences were found between sexes (Kawakami *et al*. 2014; Zhang *et al*. 2023). On the other hand, sparrows and great tits show a recombination landscape dominated by female recombination (van Oers *et al*. 2014; McAuley *et al*. 2024). However, recently there has been an effort to re-characterise heterochiasmy in crossover placement with two studies in the great reed warbler (*Acrocephalus arundinaceus*) and the sparrow (*Passer domesticus*) showing that there are heritable sex-differences in recombination in birds (Zhang *et al*. 2023; McAuley *et al*. 2024). Although fine-scale information is not available for other bird species making a comparison impossible at the moment, an emerging pattern is that descriptions of recombination patterns at the linkage group level might not reveal the whole picture of heterochiasmy in birds and a more thorough quantification of sex-specific recombination is required.

While in birds, studies are starting to quantify patterns of heterochiasmy in other species we have evidence of its existence (e.g. Kong *et al*. 2010; Brekke *et al*. 2023). However, the evolutionary reasons behind its existence remain unexplained (Burt et al., 1991; Mank, 2009; Sardell & Kirkpatrick, 2019). Identifying its causes is a complicated task since multiple factors can affect recombination in each sex (Sardell and Kirkpatrick 2019). There is evidence of molecular mechanistic processes, like the length of the synaptonemal complex (Tease and Hultén 2004; Kong *et al*. 2004; Phillips *et al*. 2015; Brick *et al*. 2018) and sex-specific standing genetic variation (Kong *et al*. 2008; Johnston *et al*. 2016; Halldorsson *et al*. 2019) that leads to heterochiasmy but a conclusive adaptive explanation for its maintenance is missing. Of the possible explanations put forward, the meiotic drive hypothesis (Brandvain and Coop 2012) suggests that in female meiosis, uncoupling a driving locus from its centromere should be favoured if drive is deleterious to the organism. Therefore, this hypothesis predicts a female increase of recombination in the region around centromeres, a pattern which appears to be most typical in empirical work (Johnston *et al*. 2017; Sardell and Kirkpatrick 2019). To our knowledge, in the avian clade, heterochiasmy has yet to be associated with centromeres and telomeres although a male-biased recombination at the ends of linkage groups has been identified (this study and Zhang et al., 2023). This lack of conclusive results can be linked to a poor genomic annotation of centromeric and telomeric sequences and future work should concentrate on properly annotating available resources before looking for the possible causes of fine-scale differences between sexes.

### Variation in recombination rate among linkage groups

Our results show that barn owl linkage groups recombine at most twice per meiosis. This result is in line with an expectation of one cross-over per chromosome (or chromosome arm) and the generally small acrocentric (or telocentric) chromosomes in the barn owl karyotype (<70Mb) (Coop and Przeworski 2007). However these recombination frequencies contrast with results from other bird species. For example, the syntenic chromosome 2 of the chicken and the flycatcher with an approximate length of 150 Mb recombines approximately six times (300 cM) (Groenen *et al*. 2009; Kawakami *et al*. 2014). In other species like the great tit and the superb fairywren, estimates match the one from our study with 100 cM for the same syntenic chromosome 2 (van Oers *et al*. 2014; Peñalba *et al*. 2020). The source of this variation in the order is unknown. Reasonable hypotheses include the localised suppression of recombination in some species (for example through segregating structural variations like inversions) or interspecific variation in the strength of crossover interference (Kirkpatrick 2010; Otto and Payseur 2019).

Some linkage groups seem to recombine less than once per meiosis (genetic length < 50 cM). This is unlikely to be the true recombination frequency of these linkage groups since the absence of an obligate crossover can lead to aneuploidy which coupled with the linkage groups’ intermediate size should generate severely deleterious consequences (Hassold *et al*. 2007). On the other hand, this observation can be due to parts of the DNA sequence missing from the assembly or filtered out during quality control, which can lead to missed distal crossovers. A larger sample size and/or a more complete assembly that incorporates the distal parts of all chromosomes might help identify the cause.

In our study, recombination rates vary substantially between and within chromosomes. As expected from an obligate crossover mentioned above, smaller chromosomes tend to have higher rates (per bp) of recombination compared to longer chromosomes. In addition, smaller chromosomes show a more uniform distribution of recombination rates along their length. On the other hand, longer chromosomes exhibit a U-shaped pattern, with reduced recombination in their centre, regardless of centromere position. In our dataset, this effect seems to diminish with increasing length implying that in the barn owl, even longer chromosomes would not further impact the skewness of recombination rates. Haenel et al. (2018) in a meta-analysis of recombination rates of different species proposed a model where the length of a chromosome and the distance from the telomere are the major factors impacting recombination rates. Their hypothesis is built upon the observation of U-shaped patterns of recombination that are present on the longer chromosomes of many species. This model was recently extended by Brazier & Glémin (2022) based on a large dataset of plant linkage maps, to include centromeric position and the placement of a single crossover per chromosome. While these advances provide an important attempt to explain broad-scale patterns guiding recombination, important questions still remain unanswered. Specifically, it is unclear if this distinction between high and low recombination regions follows a compartmentalisation of the genomic sequence into active and inactive chromatin and how these broad scale patterns define or are guided by a fine-scale hotspot landscape (Hildebrand and Dekker 2020; Jerković and Cavalli 2021). Furthermore, it is unclear if results are unique to specific classes (e.g. plants).

Even if the causes of the variation of recombination within and between chromosomes remain unexplained, some of their consequences can still be glimpsed. The most striking consequence of the unequal distribution of recombination rates along a specific length of sequence is the impact it has on nucleotide diversity. In the linkage groups studied, the Gini coefficient correlates negatively and strongly with the average nucleotide diversity. Such an outcome is expected through the action of linked selection (Begun and Aquadro 1992; Charlesworth and Jensen 2021). If recombination is spread throughout the length of the sequence, neutral alleles are uncoupled faster from selected ones which tend to drag them to extinction or fixation, thus allowing an increase of standing variation. On the contrary, long stretches of reduced recombination, through the action of linked selection, lead to reduced diversity (Charlesworth *et al*. 1993; Charlesworth and Jensen 2021). This reduced diversity can have multiple implications. It can impact observed homozygosity and affect the distribution of runs of homozygosity (ROH), as seen in other studies through a negative correlation of ROH and recombination rates (Pemberton *et al*. 2012; Bosse *et al*. 2012; Hewett *et al*. 2023). Reduced diversity can also affect estimates of divergence between populations, measures often used to identify local adaptation leading to biases and misleading inference (Charlesworth 1998; Burri 2017; Booker *et al*. 2020). Because of such implications, recombination variation can be a confounding factor in many analyses and should be accounted for when possible.

### Local and global hotspots show differences between populations

A major distinction among the recombination landscapes of species studied thus far is the presence or absence of the *PRDM9* gene. In mammals, *PRMD9* directs the recombination machinery in specific genomic regions through the generation of H3 lysine K3 tri-methylation marks (H3K4me3). In species that lack the *PRDM9* gene, including all birds, H3K4me3 marks are concentrated in regions of accessible chromatin like promoter regions of genes (Baker *et al*. 2017). Because these regions show an increased frequency of recombination, co-localise with CpG islands (CGIs) and transcription start or end sites (TSS, TES), a correlation of these genomic elements with recombination is often observed in species without *PRDM9* (Auton *et al*. 2013; Singhal *et al*. 2015; Lam and Keeney 2015; Kawakami *et al*. 2017; Baker *et al*. 2017; Schield *et al*. 2020). Our results illustrate a marked recombination increase around TSS,TES and CGIs, confirming the hypothesis that recombination is concentrated in their close proximity in the barn owl.

While recombination hotspots have been identified through the action of *PRDM9*, species in which the gene is absent do not necessarily harbour hotspots (Kaur and Rockman 2014; Smukowski Heil *et al*. 2015). The presence or absence of hotspots is usually inferred from the skewness of recombination rates along the genome (Myers *et al*. 2005), which can be summarised with the Gini coefficient. For example, in humans a Gini coefficient of 0.8 shows a marked discrepancy of recombination between the hotspot and non-hotspot regions of the genome (Myers *et al*. 2005; Kong *et al*. 2010). In *Caenorhabditis elegans,* the Gini coefficient is 0.278 and recombination is spread along the full genomic sequence (Kaur and Rockman 2014). Barn owl chromosomes harbour Gini coefficient values across the spectrum (between 0.36 and 0.7). However, a genome-wide Intermediate value inferred here (0.6), is obviously harder to place in one or the other category so to verify the presence or absence of hotspots in our dataset, we attempted to identify them. We indeed identified local hotspots heuristically, which were mostly located in regions of low recombination. Past simulation work on the power of hotspot inference shows that this is expected to be the case because power diminishes as recombination rates increase (Singhal *et al*. 2015). On the other hand, global hotspots defined following Halldorsson et al., 2019 were found in different genomic regions than local hotspots. Both classes were supported by an increase of GC content, either through the action of GC-biased gene conversion or through their co-localisation with GC-rich regions as mentioned above (Eyre-Walker 1993).

A last implication of missing the *PRDM9* gene is the evolutionary stability of recombination hotspots. The Zinc-finger domain of *PRDM9* evolves quickly changing its target sequence leading to differences in the localisation of hotspots between species and individuals that carry different alleles (Coop *et al*. 2008; Myers *et al*. 2010; Kong *et al*. 2010; Axelsson *et al*. 2012). In its absence, the functional regions that ‘attract’ the recombination machinery are hypothesised to remain stable for millions of years leading to a stable fine-scale recombination landscape (Singhal *et al*. 2015; Lam and Keeney 2015). However, the local hotspots inferred in this study showed very small overlap between pairs of populations. Even when comparing subsets of individuals from the same population (CH with CH13 and CH13_2) hotspot sharing was lower than 50%. In addition, we observed a large increase in the number of hotspots identified in populations with a smaller sample size. These observations create doubt about the biological usefulness of this definition of LD-based local hotspots in this study setting. Specifically, while few local and global hotspots coincide, and the GC content increase provides support for the existence of some local hotspots, the set identified is expected to harbour multiple false positives. Errors in hotspot inference are expected under the effects of demography (Johnston and Cutler 2012; Dapper and Payseur 2018), or simply through errors in inference (Raynaud *et al*. 2023), but in our case a sample size effect cannot be ruled out. Because the sample sizes and methods we use to infer LD-based recombination are often used for non-model species, we suggest researchers rigorously validate inferred hotspots before drawing conclusions about their evolution or stability.

Beyond hotspot sharing, the similarity of recombination landscapes was only validated at broad scales. Our studied populations diverged after the last glacial maximum when they expanded out of an Iberian refugium (Machado *et al*. 2022a; Cumer *et al*. 2022a). This timescale coupled with the dispersal abilities of the species has led to a shallow genetic differentiation (Altwegg *et al*. 2003; Machado *et al*. 2022a). The lack of a convergent fine-scale recombination landscape is not expected from a species without *PRDM9* and along with the limited hotspot sharing above, challenges the universal view of conserved recombination landscapes, as also illustrated recently in cichlid fish (Talbi *et al*. 2024). However, in our case we are cautious in interpreting such results as a true divergence of the fine-scale recombination landscape. The dependence of inference on the LD patterns and standing variation can confound results, especially in the finer scale where statistical noise increases and where even subsampled datasets can show large divergence (Raynaud *et al*. 2023; Talbi *et al*. 2024). Thus, whether this result supports a divergent fine-scale landscape or method limitations remains unclear. Future work should be cautious when using fine scale estimates in non-model species and might benefit from corroboration of results using multiple methods of inference or with verification through repeated subsampling of datasets.

## Conclusion

We present the recombination landscape of the barn owl using both linkage mapping and LD-based inference. The species is now equipped with a genome assembly with distinct linkage groups identified and a recombination map. The barn owl is thus, the first owl species with significant genomic resources paving the way for further analyses like genome-wide association studies and haplotype phasing. From our investigation of recombination rates in the species, we verify that conclusions about recombination reached in the passerine order apply to a broader phylogenetic context in the avian class. However, we caution for more conservative conclusions when using hotspots inferred through LD-based methods.

## Methods

### Samples and Sequencing

A total of 333 barn owl samples from Switzerland were sequenced for this study. In Western Switzerland breeding barn owls have been monitored for over 30 years with the installation of nest boxes. During the breeding season, the nest boxes are controlled for occupancy every 4 weeks. Individuals are ringed, measured and a blood sample is taken from their brachial vein to sex and genotype each individual to get family information (Py *et al*. 2006; Roulin *et al*. 2007; Antoniazza *et al*. 2010). Adult parents are also captured when possible and subjected to the same treatment. Using the ring identifiers of parents and offspring and the fact that barn owls show rare extra-pair paternity (Roulin *et al*. 2004), an observational pedigree has been constructed for the population.

285 individuals belonging to families from 1994 to 2020 based on pedigree information were sequenced in 2020 and 2021. Initially, we sequenced families from the pedigree that had more than 4 offspring and grandparent information whenever possible. Sample DNA was extracted from blood using DNeasyBlood & Tissue kit (Qiagen) following manufacturer’s instructions, quantified with dsDNA HS Qubit kit (ThermoFisher), diluted to 6.3 ng/μL with 10 mM Tris-HCl pH 8.0 in 40 μL. Libraries were prepared with Nextera DNA Flex (Illumina) and sequenced with Illumina Hiseq 4000 at the Lausanne Genomic Technologies Facility (GTF) We increased the dataset of sequenced individuals by including 54 more owls samples. Six originated from Georgia, and 48 from Switzerland chosen so that they had the maximum number of descendants based on the field pedigree. We sequenced these samples in 2021. Sample preparation was as above and sequencing was performed using Illumina NovaSeq 6000. All sequencing took place at the Lausanne Genomic Technologies Facility (GTF, University of Lausanne, Switzerland).

### Variant Discovery & Filtering

All available barn owl sequences were used for variant discovery. This included individuals mentioned above and samples from previous sequencing efforts (Machado *et al*. 2022a; b; Cumer *et al*. 2022a; b, 2024) along with 3 owls from the island of Corsica (Table S2 in File S1). In total, 502 samples were processed through the variant discovery pipeline described below. Raw reads were processed with trimmomatic v0.39 (Bolger *et al*. 2014). Sequence adapters were removed and reads with a length less than 70 bp were excluded. Mapping was performed with BWA-MEM v0.7.17 (Li 2013) on the barn owl genome assembly (https://www.ncbi.nlm.nih.gov/nuccore/JAEUGV000000000) (Machado *et al*. 2022a) and read groups were added with samtools v1.15.1 (Li *et al*. 2009). Since the GATK v4.2.6 (Auwera *et al*. 2013) pipeline was used for variant discovery, base quality score recalibration (BQSR) was performed using a previously published variant “truth set” (Cumer *et al*. 2022a). GATK’s Haplotype caller was run with default parameters for each individual separately to generate individual *gvcf* files.

These files were merged and joint calling was performed with all individuals together using *GenotypeGVCFs*. We initially identified 30,620,917 variants in the dataset. Filtering focused on bi-allelic SNPs and consisted of the core technical filters suggested in the GATK pipeline, a “mappability” mask and a manual individual depth filtering. Specifically, technical filters included the following criteria: QD<2.0, QUAL<30, SOR>3.0, FS>60.0, MQ<40.0, MQRankSum<-12.5 and ReadPosRankSum<-8.0. A further filtering was the exclusion of regions of the genome where our ability to confidently map reads is limited (i.e. a “mappability” mask) (Corval *et al*. 2023). Briefly, the reference genome was split into reads of 150 base pairs (bp) with a sliding of 1 bp. These artificial reads were mapped back to the reference using bwa-mem. Regions of the reference sequence where less than 90% of the reads mapped perfectly and uniquely were discarded. Variants were also filtered based on individual depth. A minimum and a maximum cutoff were applied. For the minimum cutoff, any genotype with less than five reads supporting it was set to missing (Benjelloun *et al*. 2019). For the maximum, a distribution of autosomal read depth per individual was extracted for a region (Super-Scaffold_1 and Super-Scaffold_2) with a length of 133.5Mb. The mean and standard deviation of depth was estimated and any genotype with a read depth of more than than three standard deviations from the mean was set to missing to avoid the effect of repeated regions. After filtering 26,933,469 variants were kept in 1,080 Mb of callable sequence corresponding to 1 SNP per 40 bp.

### Pedigree and relatedness

The pedigree from observational data was confirmed with genomic information from a subset of the genome. SNPs from a subset of the genome, specifically three scaffolds (Super-Scaffold_11,12, and 14) were filtered for minor allele count (>5), missing data (<10%) and were pruned for linkage disequilibrium using plink v.1.9 (Chang *et al*. 2015) with the command –-indep-pairwise 100 10 0.1. This filtering created a dataset with 91,874 SNPs. A genomic kinship matrix was calculated using the Weir & Goudet, 2017 method as implemented in hierfstat (Goudet 2005) R package. The kinship from genomic data was compared with the pedigree kinship, calculated using the kinship2 (Sinnwell *et al*. 2014) R package, and the pedigree was completed by manually resolving the first and second degree links when those could be resolved. Both k1 and k2 statistics in SNPRelate were used to discern between relationships with the same kinship value (e.g. parent-offspring and siblings). A set of unrelated individuals was selected automatically by pruning the genomic kinship table to only include individuals with a kinship of less than 0.03125. We removed individuals with multiple high-kinship links and then we prioritised individuals with higher depth of coverage. This method left a subset of 187 unrelated individuals of which 76 were from the Swiss population.

### Linkage Mapping

Lep-MAP3 (LM3) (Rastas 2017) was used to create a linkage map. A set of 250 individuals in 28 families was used, where a family in LM3 is defined as a set of individuals around a unique mating pair. Most families had 4 offspring and 4 grandparents but numbers differed and ranged from 2 to 8 offspring and from 0 to 4 grandparents. To run LM3 a stringently filtered dataset of bi-allelic SNPs was used. Specifically, we removed mendelian incompatibilities using bcftools’ mendelian plugin (Danecek *et al*. 2021) and retained a minimum of 5% MAF and a maximum of 5% missing data. We also filtered out SNPs that were less than one thousand base pairs (1 kb) apart using VCFtools (Danecek *et al*. 2011). The first step of the LM3 pipeline *ParentCall* was used to transform the data into the appropriate LM3 format and the options *halfSibs* and *removeNonInformative* were included. Data was filtered in LM3 using the *Filtering2* command, to shrink the size of the dataset. Specifically, *dataTolerance* was set to 0.01 as suggested by the author and *missingLimit* and *familyInformativeLimit* were set to 28. This meant that only variants that were non-missing and informative in all families were kept. After filtering the dataset, we retained 163,950 variants. We used *SeparateChromosomes* to identify the putative linkage groups (LGs) based on a user-defined logarithm of odds (LOD) score cutoff. We selected a LOD score of 15 (for a justification see Supplementary text – making a linkage map in File S1). Finally *OrderMarkers* with the *usePhysical* option was executed. Ordering was repeated thrice, and the output with the best Likelihood was selected for each linkage group. All three runs were compared to test for variation in estimated genetic maps. We also tested the effect of three mapping functions (Morgan’s, Haldane’s and Kosambi’s) on the estimated genetic maps (Figure S8 in File S1).

In linkage mapping certain markers might be erroneously mapped especially at the extremities of the LGs. Thus, all markers with a jump larger than 2 cM in a region of 100 markers around the ends of the LGs were filtered out. A homemade script inspired by *LepWrap* (Dimens 2022) was used. We also pruned the resulting Marey maps (plot of cM position on physical position) by regressing the genetic rank of markers with their physical rank across a linkage group. Markers with an absolute residual value of more than 100 were removed to reduce noise in the resulting maps. Finally we fitted a generalised additive model using the R Package *mgcv* (Wood 2011) and *scam* (Pya and Wood 2015) forcing a monotonically increasing smoothing spline. This makes sure that the next cM position will be larger than the previous one and gives a better fit to the data (Figure S9 in File S1).

### Linkage disequilibrium recombination

To run SMC++ (Terhorst *et al*. 2017), we followed the authors’ instructions as presented on the software’s GitHub page (https://github.com/popgenmethods/smcpp). In summary, missing data were re-coded using Plink2 (Chang *et al*. 2015) and the 5 samples with the highest coverage were selected as individuals to be provided to SMC++. The command *vcf2smc* was run for each of these five individuals. When executing ‘vcf2smc’ the mappability mask was excluded by using the –m option. The model was estimated using all output files from the previous step and with a mutation rate of 4.6e-9 estimated from family data of a collared flycatcher (Smeds *et al*. 2016). The csv-formatted estimate of piecewise-constant effective population size in past generation intervals was used in subsequent pyrho (Spence and Song 2019) analyses.

We ran pyrho with an unphased set of markers for 76 unrelated Swiss individuals. The first step in the pyrho implementation was the pre-calculation of a two-locus likelihood look-up table. This step takes into account the Ne estimates from SMC++ (Figure S2 in File S1). For the Swiss samples, the number of diploid individuals was 76 and we used the Moran approximation with a size of 200. After the inference of the lookup table the “hyperparameter” command was run to estimate metrics on the performance of different window sizes and block penalties. The authors’ guidelines were followed on how to select the best combination of parameters. Briefly we summed the Pearson correlation statistics outputted by pyrho and plotted their total sum against the L2 values. The authors suggest (https://github.com/popgenmethods/pyrho#hyperparam) that depending on the implementation one might opt to choose the parameter combination that maximises the correlation measures or minimises L2. In our case both conditions were satisfied with one combination of parameters and we run pyrho with that set of parameters. A table of the hyperparameter values for all populations can be found in Table S3 in File S1. With the inferred hyperparameters, the recombination rate was estimated using the *optimise* command on *vcfs* containing individual scaffolds which were previously filtered for singletons and a minimum distance of 10bp between variants as in Wall et al., 2022.

### Downstream analyses

After identifying linkage groups, synteny with the chicken <u>reference genome</u> was inferred with the method presented in Waters *et al*. 2021. Synteny match can be found in Figure S10 and Table S1 in File S1). The synteny includes Super-Scaffold_13 and Super-Scaffold_42 previously found to belong to the Z chromosome, as verified here (Machado *et al*. 2022a). Rate of intra-chromosomal shuffling was calculated from recombination rates inferred from mapping distances following Veller et al., 2019. Recombination rate estimates from pyrho were averaged across non-overlapping windows of different lengths using a custom script. Windows of sizes 1 kb, 10 kb, 100 kb and 1 Mb were created from the reference sequence using *bedtools makewindows* from bedtools v2.3 (Quinlan 2014). These windows were overlapped with the pyrho windows and the recombination rate in cM was calculated by multiplying the recombination probability estimate with the length of each interval and then translating this to cM using Haldane’s function (Haldane 1919). For each window, nucleotide diversity was calculated using VCFtools and the –*-window-pi* command. Estimates were corrected for masked nucleotides in each window. Sequence GC content was calculated using the reference sequence and the *bedtools nuc* command. We annotated CpG islands using the UCSC genome browser CpG island annotation tool *cpg_hl* (Kent *et al*. 2002) with default parameters. The Gini coefficient was calculated using Desctools v.0.99 (Signorell 2023). Transcription start and end sites were annotated using the genome annotation from <u>NCBI</u> as the first and last positions of the genomic sequence for each gene. Intersection of different bed files was performed using bedtools. Local hotspots were annotated by dividing the estimate of recombination rate in each focal window with the average recombination in 80 kb around (40 kb upstream and 40 kb downstream). Global hotspots were annotated as windows with at least 10 times the genome average recombination rate.

Colour palette used is *‘mako’* from the R-package viridis v.0.6.4 and is consistent throughout the figures (Garnier *et al*. 2023). Images of owls come from PhyloPic (https://www.phylopic.org). Map in Figure 4A was made using tmap v3.3-4 (Tennekes 2018) and the Natural Earth high resolution dataset. Corrplot v.0.92 was used for correlation plot in Figure 4C (Wei and Simko 2021). Vioplot v0.4.0 was used to create the violin plot in Figure 4D (Adler *et al*. 2022). Tidyverse v.2 was used for data management (Wickham *et al*. 2019). All analyses were executed in R v.4.3.1 (R Core Team 2023) using the Rstudio IDE (Posit team 2022). Light figure modification was performed in Adobe Illustrator. Scripts with commands used for data generation and downstream analyses can be found in https://github.com/topalw/Recombination_barn_owl.

## Data availability statement

Sequence data used in the study from previous publications are available on NCBI under Bioprojects codes, PRJNA700797, PRJNA727915, PRJNA727977, PRJNA774943 and PRJNA925445. Data generated for this study are available on NCBI under Bioproject code XXXXX. Code to reproduce the figures can be found in https://github.com/topalw/Recombination_barn_owl. Downstream data to be used with aforementioned scripts can be found in XXX.

## Author contributions

A.T. and J.G. devised the project. A.R. and N.P. provided the samples. A-L.D., C.S. and A.P.M. carried out the DNA-extraction and library preparation. A.T., E.L., and T.C. generated and filtered the variant dataset. A.T. performed analyses and wrote the manuscript with input from all co-authors.

## Acknowledgements

The authors would like to thank Museum national d’Histoire naturelle in Paris for the Corsican samples used in the variant discovery. They would also like to thank Julien Marquis and Melnaie Dupasquier of the GTF facility for assistance in library preparation and sequencing of the samples used in the study. This work was supported by the Swiss National Science Foundation (SNF, grant 310030_215709) to JG.

